# Profiling the Regulatory Landscape of Sialylation through miRNA Targeting of CMP- Sialic Acid Synthetase

**DOI:** 10.1101/2024.12.18.629271

**Authors:** Faezeh Jame-Chenarboo, Joseph N. Reyes, Thusini Uggalla Arachchige, Lara K. Mahal

## Abstract

Cell surface sialic acid is an important glycan modification that contributes to both normal and pathological physiology. The enzyme cytidine monophosphate N-acetylneuraminic acid synthetase (CMAS) biosynthesizes the activated sugar donor cytidine monophosphate (CMP) sialic acid, which is required for all sialylation. CMAS levels impact sialylation with corresponding biological effects. The mechanisms that regulate CMAS are relatively uncharacterized. Herein, we use a high throughput genetically encoded fluorescence assay (miRFluR) to comprehensively profile the posttranscriptional regulation of CMAS by miRNA. These small non-coding RNAs have been found to impact glycosylation. Mapping the interactions of the human miRNAome with the 3’-untranslated region of CMAS, we identified miRNA whose impact on CMAS expression was either downregulatory or upregulatory. This follows previous work from our laboratory and others showing that miRNA regulation is bidirectional. Validation of the high-throughput results confirmed our findings. We also identified the direct binding sites for 2 upregulatory and 2 downregulatory miRNAs. Functional enrichment analysis for miRNAs upregulating CMAS revealed associations with pancreatic cancer, where sialic acid metabolism and the α-2,6-sialyltransferase ST6GAL1 have been found to be important. We found that miRNA associated with the enriched signature enhanced pancreatic cell-surface α-2,6-sialylation via CMAS expression in the absence of effects on ST6GAL1. We also find overlap between the miRNA regulation of CMAS and that of previously analyzed sialyltransferases. Overall, our work points to the importance of miRNA in regulating sialylation levels in disease and add further evidence to the bidirectional nature of miRNA regulation.

## Introduction

Sialylation plays a pivotal role in various biological processes including brain development (1), B-cell maturation (2) and immune response (3), and its aberrant expression has emerged as a key factor in modulating the tumor microenvironment, (4–6). All sialyltransferases require the activated sugar donor cytidine monophosphate (CMP) sialic acid, which is biosynthesized by the enzyme cytidine monophosphate N-acetylneuraminic acid synthetase (CMAS, **Fig. 1**) (7). CMAS has been shown to play a key role in maintaining cellular sialylation levels in multiple biological contexts (5, 7–13). In breast cancer, silencing CMAS inhibits metastasis, reducing cell-surface sialic acid levels and altering the transcriptional profile of key genes involved in cancer progression (5, 9). Despite the importance of CMAS expression levels for cell-surface sialylation, there is much we do not know about its regulation.

**Figure 1.**
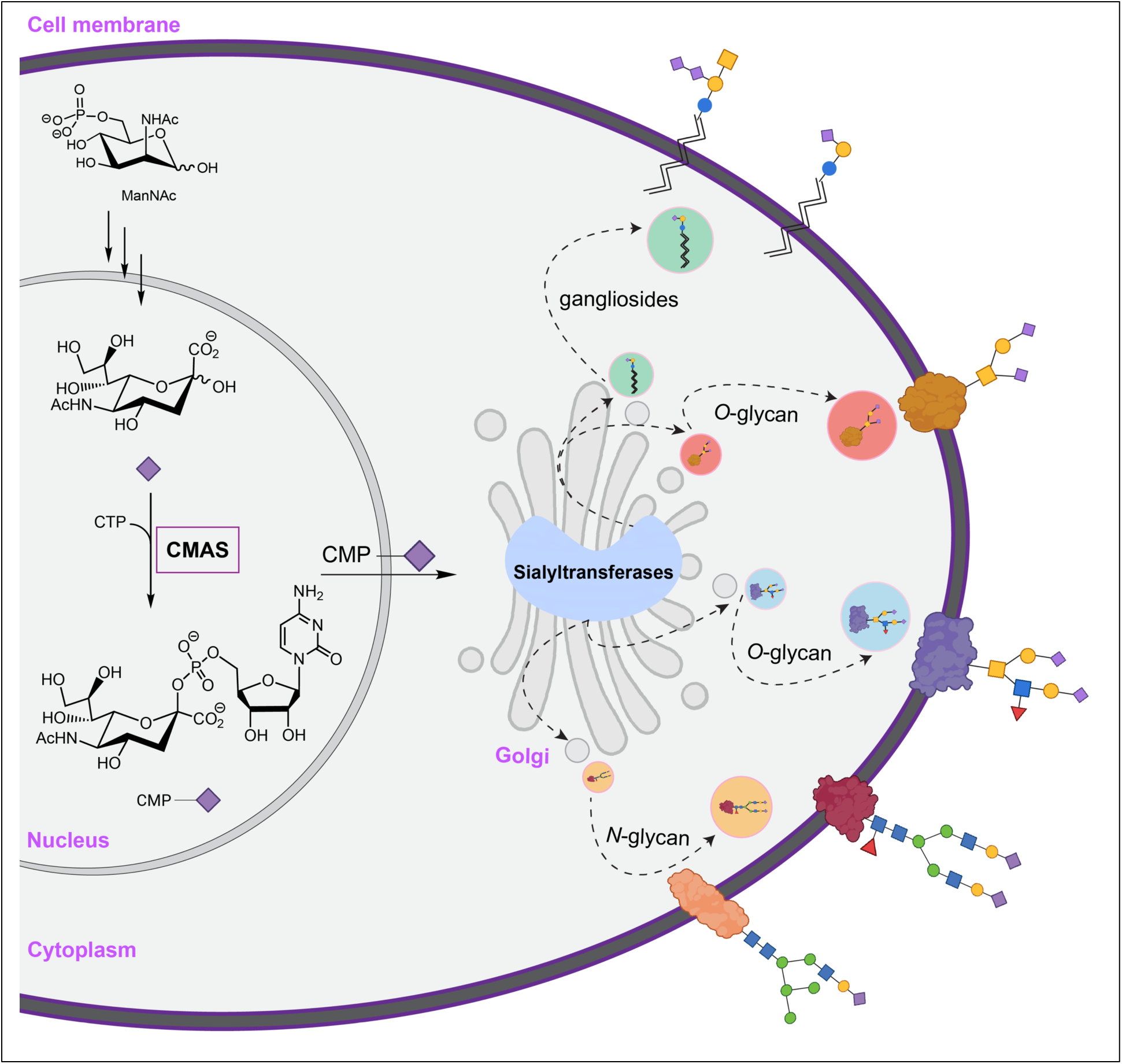
Scheme of sialoside biosynthesis. CMAS catalyzes CMP-sialic acid formation in the nucleus. It is then transported to the Golgi, where it is used by various sialyltransferases to modify glycoconjugates, including glycolipids (to make gangliosides), *N*-glycans and *O*-glycans which are transported to the cell surface. Symbolic Nomenclature for Functional Glycomics (SNFG) code is used for glycans (56). Purple diamond: sialic acid, yellow circle: galactose, yellow square: *N*-acetylgalactosamine, blue circle: glucose, blue square: *N*-acetylglucosamine, green circle: mannose, red triangle: fucose.

It is increasingly clear that posttranscriptional regulation can have strong impacts on protein expression. Our laboratory and others have shown that microRNAs (miRNAs, miRs) are important regulators of the glycome through tuning expression of biosynthetic enzymes (14–19). Most mature miRNA are ∼22 nucleotides in length and are commonly thought to inhibit protein expression by interacting with the 3’-untranslated regions (3’UTRs) of target transcripts (20, 21). Recently, our laboratory created a high-throughput assay, miRFluR, which enabled comprehensive mapping of the regulatory interactions between miRNA and a 3’UTR. Using this assay, we have mapped the miRNA interactome of 6 proteins to date, 4 sialyltransferases (ST6GAL1, ST6GAL2, ST3GAL1, ST3GAL2), a glucosyltransferase (B3GLCT) and a neutral amino acid transporter (CD98hc, aka SLC3A2)(16–18). Our analysis of these genes identified both the expected downregulatory miRNA interactions (down-miRs) and an unexpected finding of upregulatory miRNA interactions (up-miRs, i.e. miRNA:mRNA interactions that enhance protein expression) (16–18). We have shown that up-miRs, like down-miRs, act through direct binding sites between the miRNA and the 3’UTR and require Argonaute 2 (AGO2). In addition, both up- and down-miRs impact the glycome through modulation of glycosylation enzymes. miRNA are known to impact expression of multiple proteins in a regulatory network. In recent work we have shown that sialylation by ST3GAL1/2 controls the stability of CD98hc in melanoma (22) and that miRNA involved in melanoma upregulate both ST3GAL1/2 and CD98hc, providing evidence for bidirectional tuning in networks (17).

Herein, we comprehensively profiled the miRNA regulatory landscape of CMAS, screening the human miRome using our miRFluR assay. We identified both up- and down-regulatory miRNA for CMAS and validated our findings in multiple cancer cell lines. Performance of functional enrichment analysis on up-miRs for CMAS pointed towards a role for CMAS in pancreatic ductal adenocarcinoma (PDAC). We validated the impact of miRNA in pancreatic cancer cells and found that miRNA impacting CMAS expression altered α-2,6-sialylation in the absence of changes in ST6GAL1, the predominant α-2,6 sialyltransferase. We identified sites for 2 up-miRs and 2 down-miRs, confirming again that the observed miRNA regulation (both up- and down-) is via direct interactions. Overall, our analysis of miRNA-mediated regulation of CMAS has provided a comprehensive map of miRNA:CMAS interactions and pointed to the posttranscriptional co-regulation of sialyltransferases with this critical sialic acid metabolic enzyme.

## Results

### High-throughput profiling of miRNA regulation of cytidine monophosphate *N*-acetylneuraminic acid synthetase (CMAS)

In previous work, we mapped the landscape of miRNA regulation for multiple sialyltransferases (ST6GAL1, ST6GAL2, ST3GAL1, ST3GAL2) (17, 18). These enzymes, many of which are associated with cancer (22–24), all utilize CMP-sialic acid, which is synthesized by cytidine monophosphate N-acetylneuraminic acid synthetase (CMAS), a key enzyme in the sialic acid biosynthetic pathway (**Fig. 1**). CMP-sialic acid levels have been found to impact global sialylation, which in turn can regulate immune recognition, cell proliferation and migration (2, 4, 5, 11, 12, 25). However, the regulation of CMAS, and thus global sialylation, by miRNA and its intersection with miRNA regulation of specific sialyltransferases is unknown.

We profiled the miRNA regulatory landscape of CMAS using our recently created high-throughput miRFluR assay (**Fig. 2A**). This assay uses a genetically encoded dual-color fluorescence-based sensor in which the 3’UTR of a gene of interest is cloned downstream of the fluorescent protein Cerulean. The sensor plasmid also contains mCherry under an identical promoter. miRNA that either down- or up-regulate protein expression can be identified through changes in the Cerulean/mCherry ratio of the sensor. We cloned the most prevalent CMAS 3’UTR into our pFmiR sensor (**Figs. S1-2,** pFmiR-CMAS). This 3’UTR is 357 nucleotides (nts) in length, making it shorter than the median 3’UTR for both human transcripts (1200 nt) (26, 27), and the sialyltransferases studied to date (ST6GAL1: 2.75 kb, ST6GAL2: 5 kb (18), ST3GAL1: 4.9 kb, ST3GAL2: 2.25 kb (17)). We co-transfected our pFmiR-CMAS sensor with a human miRNA mimic library (Dharmacon, v. 21, 2601 miRs) arrayed in triplicate in 384-well plates into HEK293T cells. After 48 hours, the Cerulean and mCherry fluorescence was observed and the ratios were calculated. Typically, the ratiometric data is compared to non-targeting controls from the company that are found in each plate. These controls should align with the median signal in the plates. However, upon data analysis, we found that the “non-targeting” controls (NTCs) provided were significantly skewed from the median signals and shifted strongly towards downregulation across all plates (**Dataset S1**). The controls are miRNA from other organisms and their interactions with human 3’UTRs are not well studied. In previous work, we found similar interactions between the NTC and the 3’UTRs of CD98hc, ST3GAL1, and ST3GAL2. Median normalization of the datasets and validation of new controls chosen from the median miRNA allowed us to study miRNA regulation in those systems (17). We used a similar approach herein. For CMAS, each plate of miRNA was normalized to the median of the plate. miRNA with >15% variation in measurement were removed, data was log_2_ transformed and z-scored. Hits were considered as miRNA that met a z-score threshold corresponding to the 95% confidence interval. Approximately 5% of all miRNAs that passed quality control were identified as hits (73 of 1288 miRNAs, **Fig. 2B-D, Fig. S3, Dataset S1**). This percentage is in line with our data for other glycogenes (3-6% hits). We identified an even number of upregulatory (up-miRs; n=40, 55%) and downregulatory miRNAs (down-miRs; n=33, 45%) targeting CMAS. In our sialyltransferase data, the distribution of up- to down-miRs was gene specific (up:down; ST6GAL1 (76:24); ST6GAL2 (31: 69); ST3GAL1 (90: 10); ST3GAL2 (66: 34))(17, 18) with some genes showing higher prevalence of upregulation.

**Figure 2.**
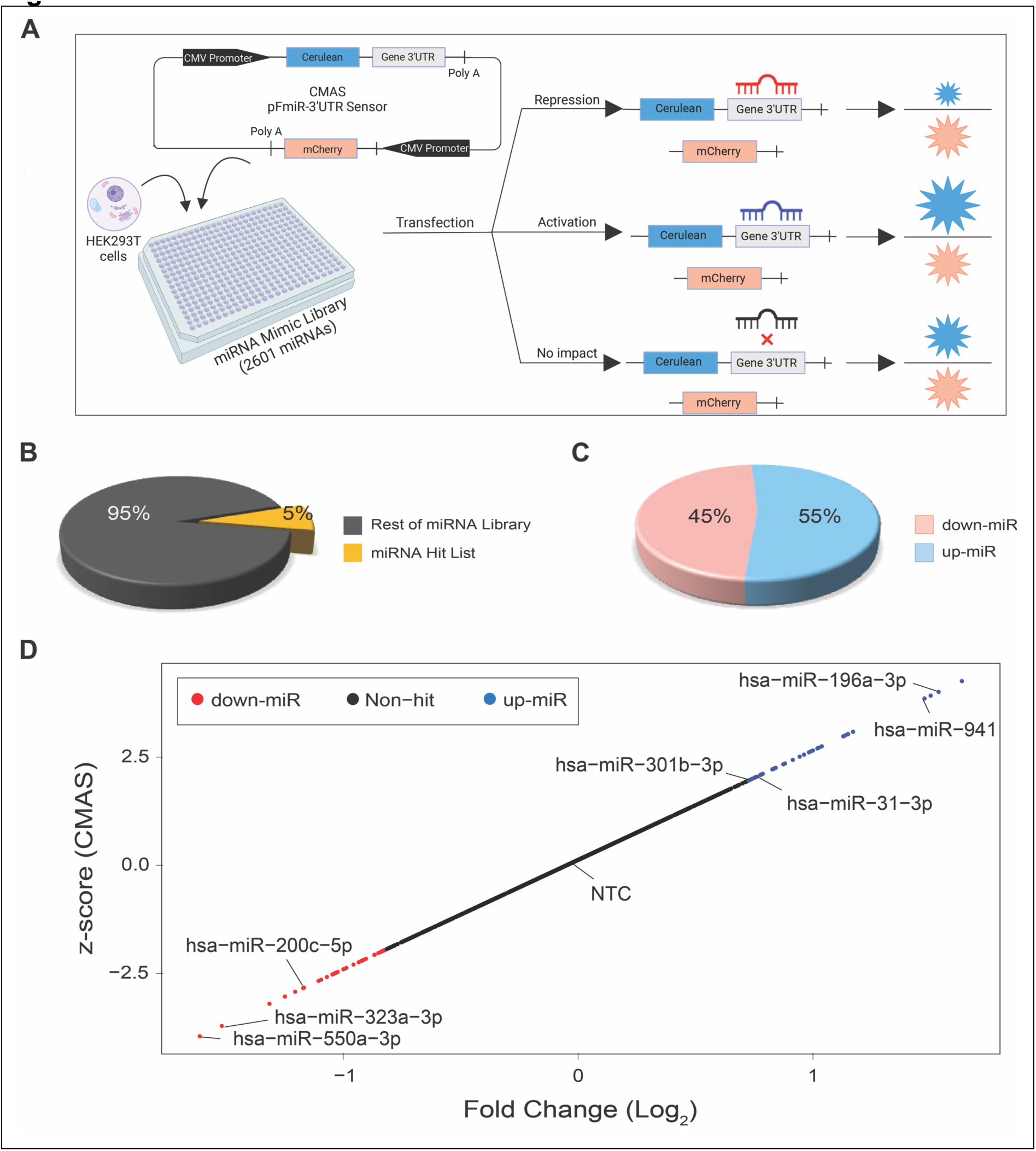
Profiling the miRNA regulatory landscape of CMAS. (A) Schematic of the miRFluR assay. pFmiR-3’UTR CMAS sensor is co-transfected with human miRNAome library and HEK293T cells in 384-well plates. (B) Pie chart comparing miRNA hits (yellow, 5%) and non-hits (grey) for CMAS. (C) Pie chart comparing percent of down-miRs (red,45%) and up-miRs (blue, 55%) in CMAS hit list. (D) Scatter plot of miRFluR data for CMAS; miRNA in the 95% confidence interval for CMAS regulation are colored (down-miRs: red, up-miRs: blue) and the tested miRNAs are labelled. All miRNA data show is post-QC.

### miRNAs bidirectionally regulate CMAS expression in breast cancer cells

Sialylation plays important roles in the biology of breast cancer promoting metastasis and shielding cells from immune recognition (5, 9, 11). CMAS expression is negatively correlated with survival in breast cancer patients (5). Knockdown of this enzyme inhibits progression and metastasis in multiple mouse models (5, 9). We thus chose the breast cancer cell lines MDA-MB-231 and MDA-MB-436 for our initial validation of miRNA regulation of CMAS. We first needed to define a new non-targeting control as that supplied by the company had a profound impact on CMAS expression in both Western blot and miRFluR assays (**Dataset S1**, **Fig. S4**). We tested two miRNAs from the median data from our array (miRs-625-5p and -548ay-3p). These miRNAs were chosen as they had few known biological interactions and thus were unlikely to have profound impacts on the cells. Neither miRNA impacted CMAS expression, in line with our other data (**Fig. S4**). We chose to use miR-625-5p as our new non-targeting control (NTC). To validate the identified miRNA hits for CMAS, we chose a subset of miRNAs previously shown to play roles in breast cancer (28, 29): down-miRs (miRs: -200c-5p, - 323a-3p, -550a-3p) and up-miRs (miRs: -301b-3p, -31-3p, -196a-3p, -941) for their impacts on endogenous CMAS expression in MDA-MB-231 and MDA-MB-436 cell lines (**Fig. 3, Fig. S5**). Overall, the miRNA regulation of CMAS expression in both cell lines was consistent with our miRFluR data. Significant results were observed in at least one cell line for all miRNA, in line with our miRFluR assay with up-miRs causing a ∼30-70% increase in protein expression and down-miRs inhibiting protein expression by ∼30-50% (**Fig. 3A-B, Fig. S5A-D, Table S1**).

**Figure 3.**
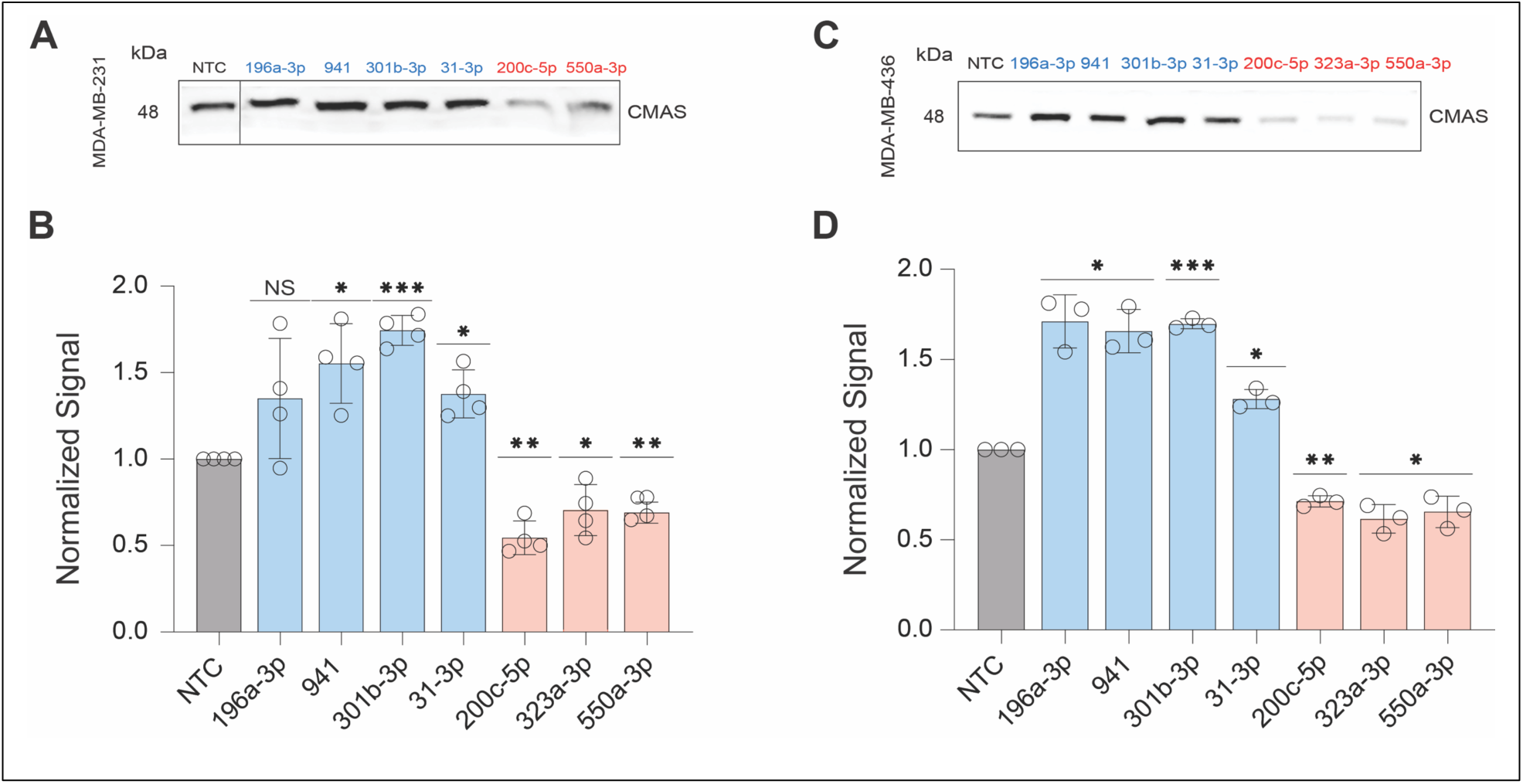
miRNA regulation of CMAS expression in breast cancer cell lines MDA-MB-231 and MDA-MB-436. (A, C) Representative Western blot analysis of CMAS in MDA-MB-231 (A), MDA-MB-436 (C). miRNA mimics (non-targeting control (NTC, black/grey): miR-625-5p which is a median control, down-miRs (red): -200c-5p, -323a-3p, -550a-3p; up-miRs (blue): -196a-3p, -941, -301b-3p, -31-3p) were transfected into cells. Western blot analysis was conducted 48 hours post transfection. (B, D) Bar charts show three independent biological replicates of analysis shown in A & C for MDA-MB-231 (B) and MDA-MB-436 (D). All Western blot samples were normalized to total protein staining (Ponceau) and then to normalized NTC. All experiments were performed in ≥ biological triplicate, individual replicates are shown. Errors shown are standard deviations. Western blot analysis: one-sample *t*-test was used to compare Ponceau normalized data for miRs to NTC. (NS: not significant, *\* p < 0.05, ** < 0.01, *** < 0.001*). The Western blot results were also analyzed using paired *t*-tests against NTC and *p*-values are reported in Table S1.

We next tested whether endogenous miRNA impact CMAS levels using miRNA hairpin inhibitors (anti-miRs) (**Fig. 4A**). In MDA-MB-231 miRs: -301b-3p, -941, -31-3p have high endogenous expression, thus we focused on these miRs. Because the anti-miR controls bind the miRNA in the Ago2 complex, the commercial negative control does not impact protein expression and thus the supplied anti-NTC was used. Upon transfection of the anti-up-miRs (anti-301b-3p, -941, and -31-3p) in MDA-MB-231 we observed a significant loss of CMAS (∼40% decrease, **Fig. 4B-C, Fig. S6, Table S1**). This strongly argues that endogenous miRNA upregulate this crucial metabolic enzyme.

**Figure 4.**
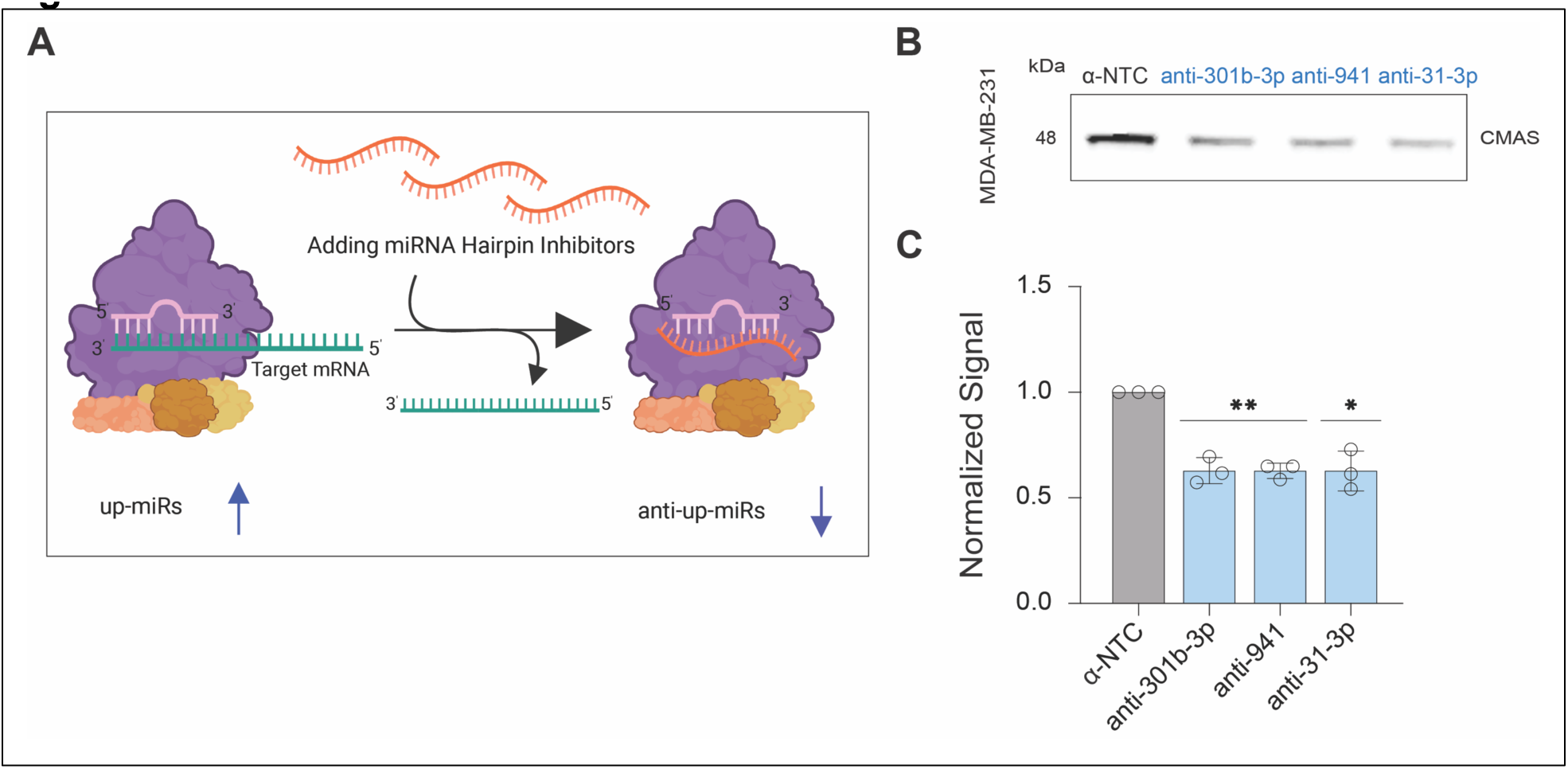
Inhibition of endogenous CMAS up-miRs by hairpin inhibitors in MDA-MB-231. (A) Schematic representation of miRNA hairpin inhibitor (anti-miR) mode of action. (B) Representative blot for anti-miR non-targeting control (α-NTC from Dharmacon) and anti-up-miRs (anti-miR-301b-3p, anti-941, and anti-31-3p) in MDA-MB-231 cells. (C) Graph of three biological replicates of analysis shown in (B). All Western blot samples were normalized to total protein staining (Ponceau) and then to normalized α-NTC. Errors shown are standard deviations. Western blot analysis: one-sample *t*-test was used to compare Ponceau normalized data for miRNAs to α-NTC, *\* p < 0.05, ** < 0.01*). The Western blot results were also analyzed using the paired *t*-test and reported in Table S1.

### Sialylation is regulated by miRNA in pancreatic cancer via modulation of CMAS expression

With the validation of our miRFluR results in hand, we next examined whether miRNAs that upregulate CMAS expression might show enrichment in specific pathways and diseases using the miRNA enrichment and annotation tool (miEAA) created by Keller and colleagues (30, 31) (**Fig. 5A-B**). Unlike other methods that use gene enrichment analysis of the targets of the miRNA, this tool uses annotations of the miRNA themselves curated from a variety of sources including literature, enabling a less biased enrichment analysis at the level of the miRNA. We performed pathway enrichment analysis on our 40 up-miRs, analyzing only pathways that had a minimum of 5 miRNA hits per subcategory. Our results indicated a role for CMAS in cancer and in pancreatic cancer specifically (**Fig. 5A, Dataset S2**). Of note, the majority of our upregulatory miRNA were implicated in cancer pathways (∼82%) and over half (∼70%) were implicated in pathways specific to pancreatic cancer (**Fig. 5A, Dataset S2**). Examination of the disease states associated with our up-miR hit list in miEAA also identified pancreatic cancer as a top hit (**Fig. 5B, Dataset S2**). We next analyzed *cmas* transcript expression in three cancers (breast (BRCA), thyroid (THCA) and pancreatic (PAAD), **Fig. 5C**) using the GEPIA web server, which contains RNA sequencing dataset for a total of 9,736 tumors and 8,587 normal samples from the TCGA database and the GTEX projects (32, 33). The result indicates higher levels of *cmas* expression in both breast and pancreatic cancers, with a significant enhancement in pancreatic tumors.

**Figure 5.**
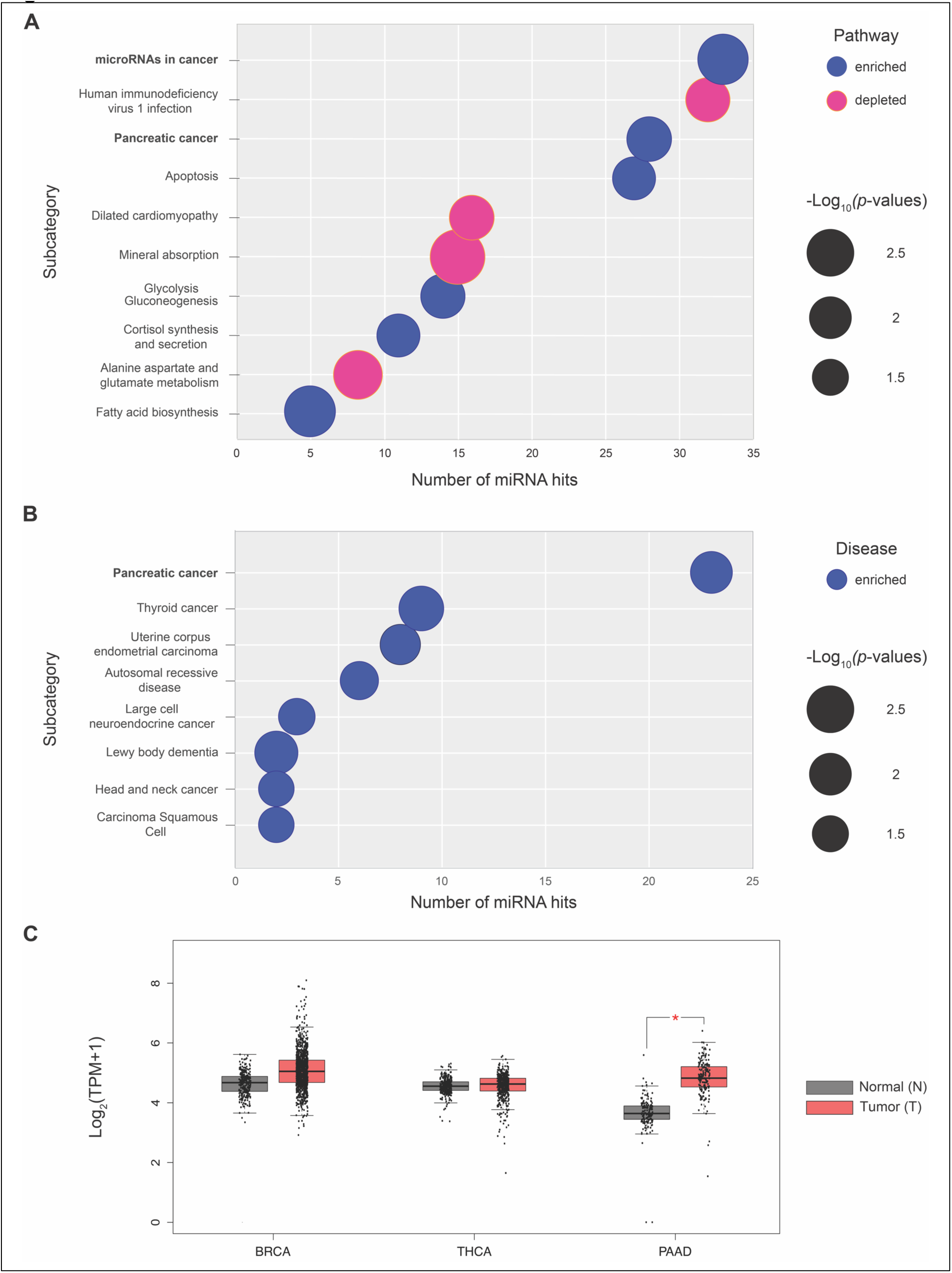
Pathway and disease enrichment analysis for miRNA upregulators of CMAS. (A) Bubble plot indicates pathways enriched in CMAS up-miRs. Blue circles indicate miRNAs are higher in pathway, while pink circles indicate miRNA are depleted in the pathway. Size of the circle indicates relative *p*-values. (B) Bubble plot of diseases enriched in CMAS up-miRs. All enrichment analysis was conducted using the miRNA enrichment analysis and annotation tool (miEAA) (31). (C) Graph of *cmas* transcript expression in normal (grey, N) and tumor (red, T) samples of breast (BRCA, N=291, T=1085), thyroid (THCA, N=337, T=512) and pancreatic (PAAD, N=171, T=179) cancers based on TCGA and GTEx datasets. Statistical analysis was done by the GEPIA database (32, 33) using one-way ANOVA. *\* p < 0.01*.

Although a role for CMAS has not been previously observed, we and others have shown both specific roles for and a strong association of sialylation with pancreatic ductal adenocarcinoma (PDAC, PAAD) (6, 34, 35). To test whether up-miRs impact regulation of sialic acid in pancreatic cancer cells, we tested one down-miR (-200c-5p), and two up-miRs (-31-3p and -196a-3p) in SU.86.86, a pancreatic cancer cell line with high levels of both α-2,3 and α-2,6 sialylation (10) (**Fig. 6B-C, Fig. S7A-B**). The up-miRs chosen both contributed to the pancreatic cancer signatures in our enrichment analysis **(Fig. 5A-B, Dataset S2).** In line with our previous results, we observed inhibition of CMAS expression by down-miR-200c-5p, whereas both up-miRs tested significantly enhanced upregulation of the enzyme in this cell line (**Fig. 6B-C**).

**Figure 6.**
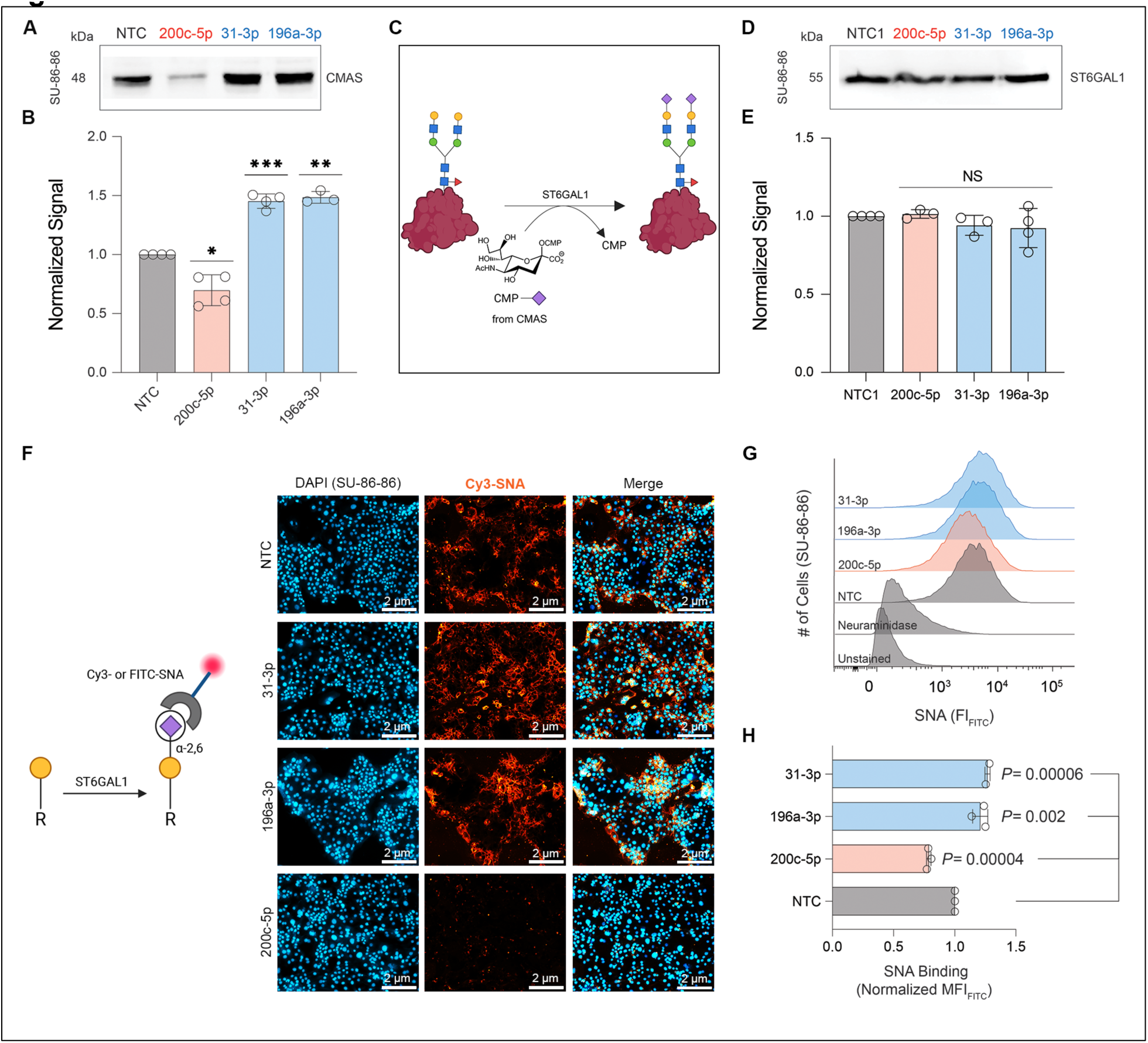
miRNA regulation of CMAS expression in SU.86.86 pancreatic cancer cell line. (A) Representative Western blot analysis of CMAS in SU.86.86 cells. miRNA mimics (down-miR (red): -200c-5p; up-miRs (blue): -196a-3p, -31-3p) or median control: miR-625-5p (NTC) were transfected into SU.86.86 cells. Cells were lysed and analyzed 48 hours post-transfection. (B) Bar chart shows three independent biological replicates of analysis shown in A. (C) Schematic representation of α-2,6-sialylation by ST6GAL1. (D) Representative blot of Western blot analysis for ST6GAL1 for cells treated as in (A). (E) Bar graph represents normalized data for biological replicates of Western blot analysis as in D. (F) Left: Schematic representation of SNA lectin binding. Right: SNA staining of cells treated with miRNA (up-miRs: -31-3p, -196a-3p; down-miR-200c-5p or median control) as in A. (G) Representative flow cytometry histograms quantifying SNA binding of cells treated as in A. (H) Bar graph of flow cytometry quantification. All experiments were performed in ≥ biological triplicate. All Western blot samples were normalized to total protein staining (Ponceau) and then to normalized NTC. Western blot analysis: one-sample *t*-test was used to compare Ponceau normalized data for miRNAs to α-NTC, NS: not significant, * *p < 0.05, ** < 0.01, *** < 0.001*).). The Western blot results were also analyzed using the paired *t*-test and reported in Table S1. For flow cytometry analysis, the paired *t*-test was used to compare miRs to NTC and *p*-values are indicated on the graph.

We next wanted to observe whether impacting CMAS would impact sialylation. Previous work identified α-2,6-sialic acid and its underlying sialyltransferase, β-galactoside α-2,6-sialyltransferase1 (ST6GAL1), as important in PDAC formation and metastasis (6, 34, 35) (**Fig. 6A**). We wanted to know whether miRNA regulation of CMAS might contribute to the high levels of α-2,6-sialylation observed on pancreatic cancers independently of regulation of ST6GAL1. None of the three miRNAs tested were observed to directly regulate ST6GAL1 in our previous miRNA analysis(18). To test for indirect impacts of these three miRNAs on ST6GAL1 levels (and thus α-2,6-sialylation), we checked the protein expression levels of ST6GAL1 upon miRNA mimic transfection in SU.86.86 cells (**Fig. 6D-E, Fig. S7C-D**). The data showed no impact of these miRNA on ST6GAL1 expression, in line with our previous study (18). We next tested whether these miRNAs impacted α-2,6-sialylation using both fluorescence microscopy and flow cytometry with Cy3-conjugated and FITC-conjugated *Sambucus Nigra* lectin (SNA), respectively, which is specific to α-2,6 sialic acid (36, 37) (**Figs. 6F, G-H**, respectively). We observed modulation of α-2,6-sialylation in SU.86.86 cells in line with the impacts of the miRNAs on CMAS levels. Together, our data provides evidence of miRNA regulation of cellular α-2,6-sialylation in pancreatic cancer by alteration in CMAS levels independent of ST6GAL1 regulation. This confirms a role for both sialyltransferase expression and metabolic flux in the hypersialylation observed in PDAC (6, 10, 34).

### miRNA regulation of CMAS expression is via direct miRNA: 3’UTR interactions

Downregulation of protein expression by miRNA is the canonical mode of miRNA action and generally requires base-pairing to the mRNA of at least 6 nucleotides at positions 2-6 at the 5’ end of the miRNA (seed region (20, 21)). In previous work, our laboratory showed that direct miRNA:3’UTR binding is also required for upregulation, although in most cases the binding pattern did not follow canonical seed binding rules (17, 18). We set out to identify the binding sites for 2 validated down-miRs (miRs-200c-5p, -550a-3p) and 2 validated up-miRs (miRs-31-3p, -301b-3p). We focused on miRNAs that were important in either pancreatic cancer (miRs: -200c-5p (38), -31-3p (**Fig. 5A-B, Dataset S2**) or breast cancer (miRs: -550a-3p (39), -301b-3p (28)). To identify down-miR sites, we used Targetscan (20, 40) and for up-miRs we used RNAhybrid (40). As anticipated, the predicted sites for downregulation showed canonical seed binding, while those for upregulation did not (**Fig. 7 A-D**). Of note, the binding sites for miR-200c-5p and miR-550a-3p were found to be identical within the 3’UTR. We then mutated the predicted sites in the CMAS 3’UTR to the sequence of the corresponding miRNA on the pFmiR-CMAS plasmid. We co-transfected the wildtype (WT) or mutant sensors with the NTC or appropriate miR mimic into HEK293T cells. For each sensor, mimic data was normalized to NTC and results of 3 biological replicates are shown in **Fig. 7E**. In all cases, the impact of the miRNA on sensor expression was lost upon mutation. This provides strong evidence that both down- and up-regulation of CMAS expression by miRNA is through direct interactions with the CMAS 3’UTR.

**Figure 7.**
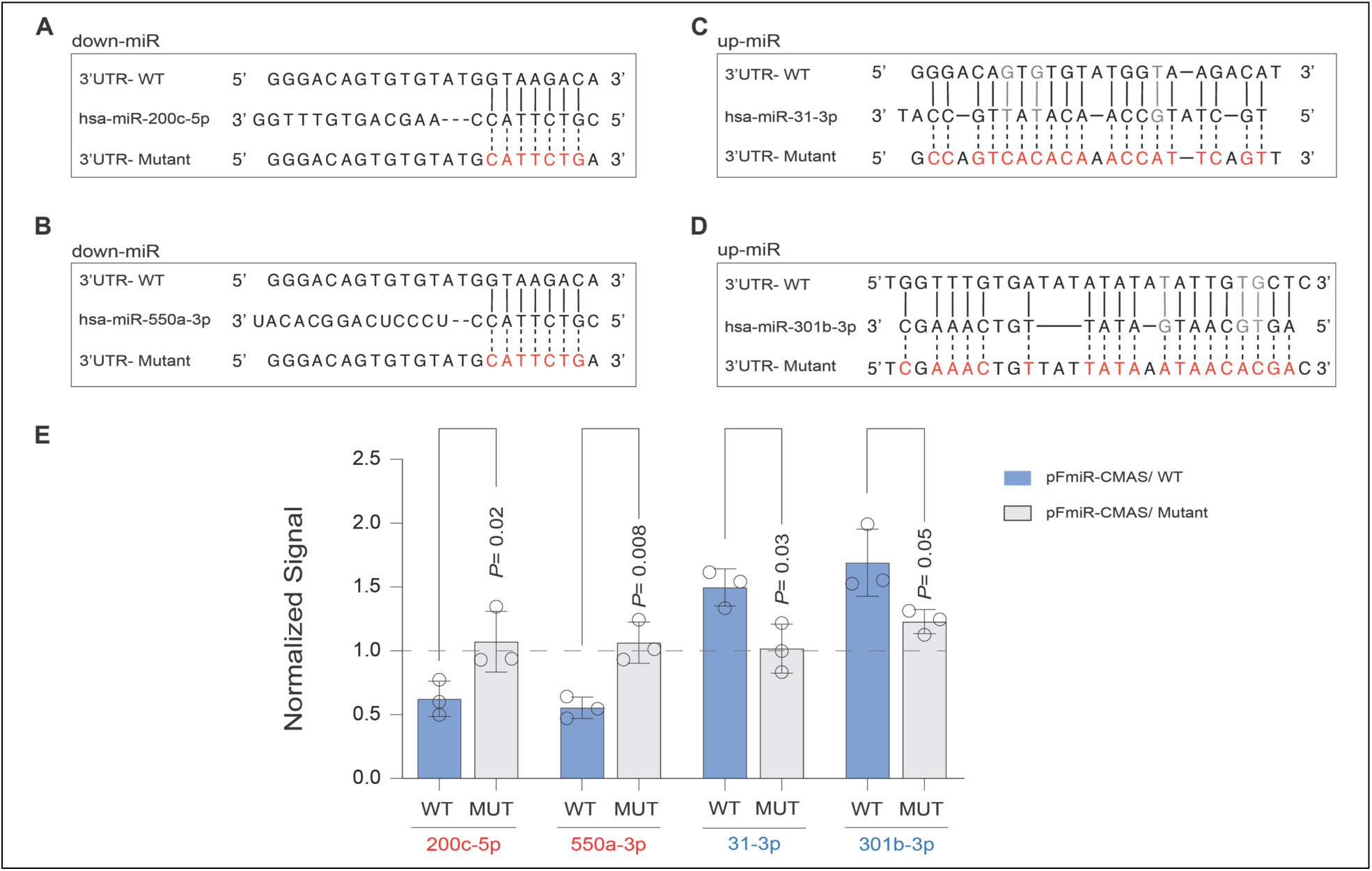
miRNA tune CMAS expression via direct interactions with its 3’UTR. (A-D) Alignment of down-miRs (A: miR-200c-5p; B: -550a-3p) and up-miRs (C: miR-31-3p; D: -301b-3p) with their predicted CMAS-3′UTR sites and the corresponding mutants. Mutated residues are shown in red, wobble interactions (G: U) are indicated in grey. (E) Bar graph of data from wildtype (WT, blue) and mutant (MUT, grey) miRFluR sensors. For each sensor, data was normalized over NTC. The experiment was performed in biological triplicate and the paired *t-*test was used for comparison. *P-*values are indicated on graph.

## Discussion

Regulation of sialylation through changes in CMAS levels affects both normal (brain development, immune response) and pathological states (cancer progression, autoimmunity) (2, 4, 5, 7, 9, 10). microRNA are a key posttranscriptional regulator of both the proteome and by extension the glycome (15–18, 41, 42). The canonical view of miRNA is that they inhibit protein expression via mRNA degradation or loss of mRNA translation (20, 21). Recently, we have shown that miRNA can upregulate protein expression through direct interactions with the 3’UTR of mRNA in actively dividing cells (16–18), a finding that has been confirmed by others (43). In line with our previous discovery, upon mapping the comprehensive miRNA regulatory landscape of CMAS, we found miRNA regulation to be bidirectional, with 33 downregulatory miRNA (down-miR) and 40 upregulatory miRNA identified by our high-throughput miRFluR assay (**Fig. 2**). The relevance of hits to regulation of CMAS levels was validated in several cell lines (**Figs. 3 & 6**) confirming both up- and downregulation of this key sialic acid biosynthetic enzyme by miRNAs.

Mutagenesis analysis of the binding sites for four of our miRNA hits (down-miRs: - 200c-5p, -550a-3p, up-miRs: -31-3p, -301b-3p) verified that both up- and downregulation by these miRNAs were through direct miRNA:mRNA interactions. The two down-miRs shared the same 8-mer seed, a canonical miRNA:mRNA binding pattern. In contrast, neither of the two up-miRs had canonical miRNA binding sites. This is consistent with earlier work, in which only 2 of the 10 up-miR binding sites validated prior to this publication were found to follow canonical seed pairing rules (17, 18, 43). Similar to our previous findings, upregulatory sites for CMAS were not AU rich, contrasting with the requirements for upregulation observed in non-dividing cells by Steitz, Vasudevan and others (44–46). Previous work found that upregulation requires Argonaute 2 (AGO2) (18, 43, 45–47). A significant percentage (∼60%) of 3’UTR:miRNA:AGO2 complexes identified by cross-linking contained non-canonical binding motifs, pointing to the common nature of such interactions (20, 48).

Regulation by miRNA controls the expression of a network of genes with a specific phenotypic outcome (14, 15, 49), which is strongly illustrated by recent work showing that miR-193 is responsible for the classic color variations in moths seen in evolution (50). In earlier work, our laboratory demonstrated that downregulation of any of the glycosylation enzymes B3GLCT, ST3GAL5 or ST6GALNAC5, targets of miR-200b-3p – a critical regulator of epithelial-to-mesenchymal transition, induce an epithelial phenotype in line with the known impact of the miRNA. Based on these findings, we proposed the miRNA proxy hypothesis which states, “miRs can be used to identify (by proxy) the biological functions of specific …proteins.”(41). In line with our hypothesis, enrichment analysis of the gene targets and disease associations of downregulatory miRNAs that target the glycosyltransferase B3GLCT identified symptoms of Peters’ Plus syndrome, a congenital disorder caused by the loss of this enzyme (16). Whether upregulatory miRNA would also be predictive was untested. Our miRNA enrichment analysis of upregulatory miRNA impacting CMAS were enriched in cancer pathways. In specific, pancreatic ductal adenocarcinoma (PDAC, pancreatic cancer) was strongly associated with the upregulatory miRNA, arguing that CMAS (and CMP-sialic acid levels) might be important in PDAC. In line with this, we observed differential *cmas* transcript levels in PDAC (**Fig. 5C**). Previous work by Rodriguez *et al,* which found that the CMP-sialic acid biosynthetic pathway as a whole was upregulated in PDAC (10), further confirms the findings of our enrichment analysis. In concordance with these expression levels, both our laboratory, and others, have shown high levels of sialylation, especially α-2,6-sialic acid, in PDAC (6, 34, 35). We found that miRNA could impact α-2,6-sialylation levels through CMAS without altering ST6GAL1, the main α-2,6-sialyltransferase in pancreatic cancer. Taken together this work shows that upregulatory miRNA also predict functional roles for the genes they target.

Canonically, miRNA in regulatory networks are presumed only to downregulate protein expression (51). We have recently shown upregulatory miRNA regulate proteins that are in functional networks, making miRNA, like transcription factors, bidirectional regulators (17). Given the role of CMAS in biosynthesizing the sugar donor for the sialyltransferases, we compared the CMAS miRNA dataset gathered herein to the miRNA datasets obtained for previously studied sialyltransferases (ST6GAL1, ST6GAL2, ST3GAL1, and ST3GAL2). We found several regulatory miRNAs in common between these transcripts (17, 18) (**Fig. 8**). Co-regulation of CMAS with sialyltransferases was in the same direction. For example, miR-217 upregulates both CMAS and ST3GAL1, while miR-363-5p downregulates both. Given the relative roles of CMAS and sialyltransferases, it makes sense that tuning would be in the same direction. Of note, miR-212-5p, which is highly expressed in PDAC, co-upregulates CMAS, ST6GAL1 and ST3GAL2 in our assays (52, 53). All three genes are upregulated in PDAC, and high levels of sialylation are observed in that cancer (6, 10, 34, 35, 54). miR-301b-3p, which we validated in this study, was a co-up-miR for both CMAS and ST6GAL2. Our miRNA enrichment analysis identified thyroid cancer as another disease associated with our CMAS upregulators, with miR-301b-3p as part of the signature (**Fig. 5B**). The upregulation of ST6GAL2 causes tumorigenesis in follicular thyroid carcinoma via impairing the Hippo signaling pathway (24). This suggests potential co-upregulation of ST6GAL2 and CMAS expression in thyroid cancer. Overall, our data supports a paradigm in which bidirectional tuning of protein expression by miRNA regulate phenotypes, including sialylation.

**Figure 8.**
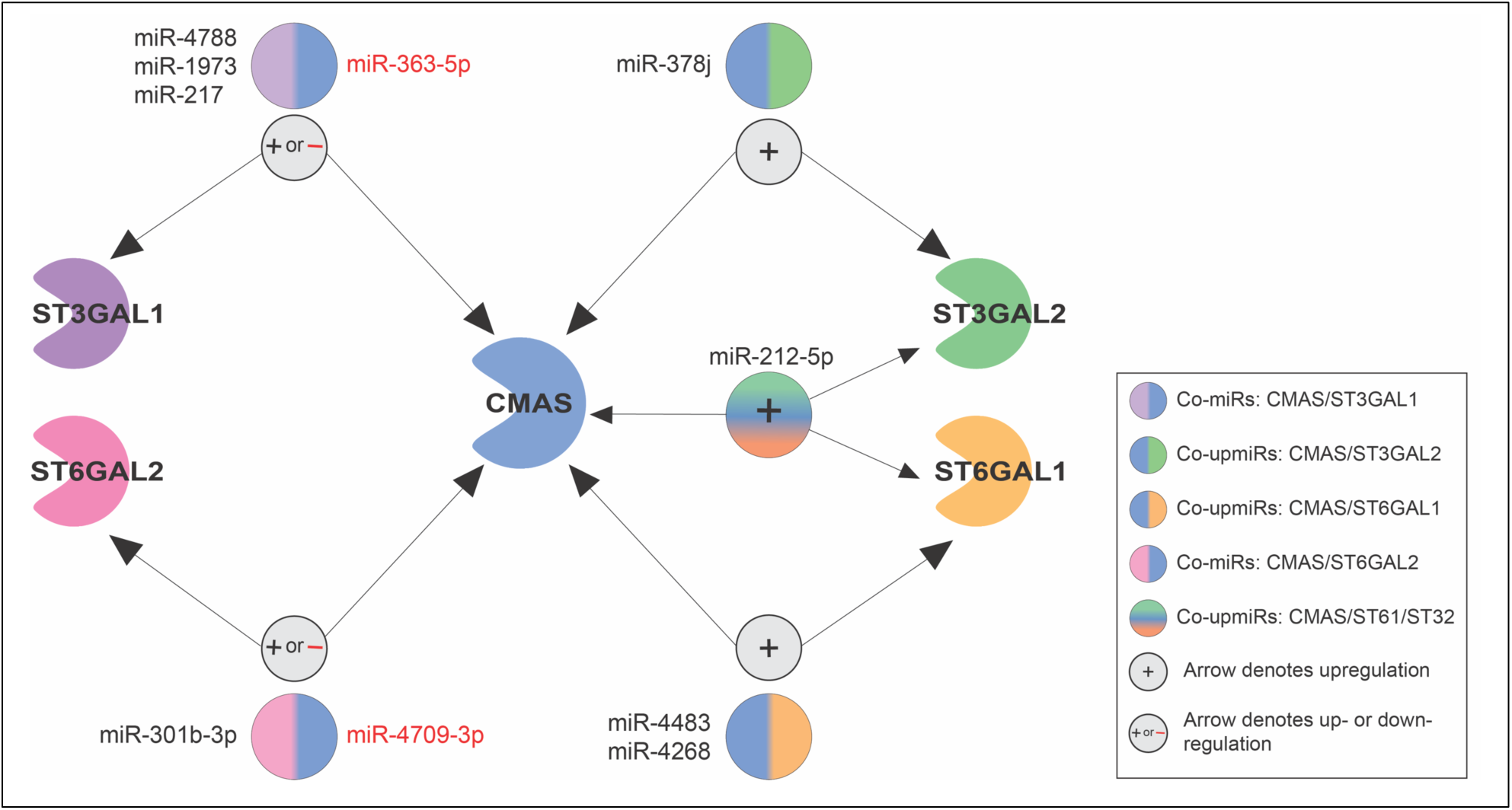
Map of miRNAs regulating CMAS and sialyltransferases. Data for miRNA regulation of 4 sialyltransferases (ST3GAL1-purple, ST3GAL2-green, ST6GAL1-yellow, ST6GAL2-pink) was overlaid with that of CMAS and co-regulators identified. miRNA are shown next to multicolored circles indicating proteins co-regulated by them. All co-regulation was either upregulatory (+, black miRNAs) or downregulatory (-, red miRNAs) and no opposing regulation was observed.

## Experimental procedures

### Cloning

CMAS 3’UTR was amplified via PCR from genomic DNA (gDNA, 10 µg, from HEK293T cell line) using the primers shown in **Table S2**. The amplicons were cleaned up using a Monarch^®^ PCR & DNA cleanup kit (catalog #: T1030S, NEB). The 3’UTR fragments were then cloned into restriction sites (NheI and BamHI) downstream of Cerulean in the pFmiR-empty backbone (16) using standard ligation protocols (NEB). Plasmids were verified by Sanger sequencing (Molecular Biology Services Unit, University of Alberta). Large-scale endotoxin free DNA preparations were made for sequence-verified constructs (pFmiR-CMAS) using Endotoxin-free plasmid DNA purification (Takara Bio USA, Inc., catalog #: 740548). The plasmid map for pFmiR-CMAS and its corresponding 3’UTR sequence can be found in **Fig. S1.**

### Cell lines

All cells lines (HEK-293T, MDA-MB-231, MDA-MB-436, SU.86.86) were purchased directly from the American Type Culture Collection (ATCC) and cultured using suggested media (HEK-293T & MDA-MB-231: Dulbecco’s Modified Eagle Medium (DMEM), 10% FBS; MDA-MB-436: DMEM, 10% FBS, sodium pyruvate; SU.86.86: RPMI-1640, 10% FBS) under standard conditions (5% CO_2_, 37℃).

### miRFluR High-throughput assay

The Human miRIDIAN miRNA mimic Library 21.0 (Horizon Discovery (*formerly Dharmacon)*) was resuspended in ultrapure nuclease-free water (REF. #: 10977-015, Invitrogen) and aliquoted into black 384-well, clear optical bottom tissue-culture treated plates (Nunc). Each plate contained three replicate wells of each miRNA in that plate (2 pmol/well). In addition, each plate contained a minimum of 6 wells containing non-targeting controls (NTC1, cat. #: CN-001000-01, Horizon Discovery). To each well was added target pFmiR plasmid (pFmiR-CMAS: 30 ng) in 5 µl Opti-MEM (Gibco) and 5 µl of transfection solution: 0.1 µL Lipofectamine™ 2000 (cat. #: 11668500, Life Technologies) diluted to 5 µl total volume with Opti-MEM (Gibco) and premixed 5 min at room temperature. The mixture was allowed to incubate at room temperature in the plate for 20 min. HEK293T cells (25 µl per well, 400 cells/ µl in phenol red free DMEM supplemented with 10 % FBS and Pen/Strep) were then added to the plate. Plates were incubated at 37℃, 5% CO_2_. After 48 hours, the fluorescence signals of Cerulean (excitation: 433 nm; emission: 475 nm) and mCherry (excitation: 587 nm; emission: 610 nm) were measured using the clear bottom read option (SYNERGY H1, BioTek, Gen 5 software, version 3.08.01).

### Data analysis

We calculated the ratio of Cerulean fluorescence over mCherry fluorescence (Cer/mCh) for each well in each plate. For each miRNA, triplicate values of the ratios were averaged, and the standard deviation (S.D.) obtained. We calculated % error of measurement for each miRNA (100 × S.D./mean). As a quality control measurement (QC), we removed any plates or miRNAs that had high errors in the measurement (median error ±2 S.D. across all plates) and/or a high median error of measurement for the plate (>15%). After QC, we obtained data for 1288 miRNAs for CMAS out of the 2601 total miRNAs screened against its 3’UTR. The Cer/mCh ratio for each miRNA was then normalized to the median of Cer/mCh ratios within that plate and error was propagated. Data from all plates were then combined and log-transformed z-scores calculated. A z-score of ±1.965, corresponding to a *two*-tailed *p*-value of 0.05, was used as a threshold for significance. Post-analysis we identified 73 miRNA hits for CMAS within 95% confidence interval (see **Fig. 2B-D**, **Fig. S2**, and **Dataset S1**).

### Western blots

Western blot analysis was conducted for CMAS in two cell lines: MDA-MB-231, MDA-MB-436 cell lines (down-miRs: miR-200c-5p, miR-550a-3p, miR-323a-3p; up-miRs: miR-941, miR-196a-3p, miR-301b-3p, miR-31-3p). A small subset of miRNA: one down-miR and two up-miRs was tested in SU.86.86 cell line: down-miR-200c-5p; up-miRs: -31-3p & -196a-3p. For Western blot analysis, miR-625-5p was selected as the non-targeting control (NTC) based on its lack of impact on CMAS in either the miRFluR assay or in cells (**Dataset S1, Fig. S4**). For all experiments, cells were seeded in six-well plates (50,000 cells/well) and cultured for 24 h in appropriate media. Cells were then washed with Hanks buffered salt solution (HBSS, Gibco) and transfected with miRNA mimics (50 nM mimic, Dharmacon, Horizon Discovery, 5 μl Lipofectamine 2000, Life Technologies in 250 μl OptiMEM). The media was changed to standard media 12 h post-transfection. Cells were then lysed at 48 h post-transfection in cold RIPA buffer supplemented with protease inhibitors (Thermofisher, catalog #: 89900). For Western blot analysis, 30 µg of protein was added to 4x loading buffer with DTT (1 mM), heated at 95 °C for 10 min and run on a 10% gel (SDS-PAGE) using standard conditions. Proteins were then transferred from the gel to Polyvinylidene fluoride membrane using iBlot2 Transfer Stacks (PVDF, Invitrogen, catalog number: IB24002) and the iBlot2 transfer device (Invitrogen) using the standard protocol (P_0_). Blots were then incubated with Ponceau S Solution (Boston BioProdcuts, catalog #ST-180) for 3 min and the total protein levels were imaged using the protein gel mode (Azure 600, Azure Biosystems Inc.). Blots were blocked with 5% BSA in TBST buffer (TBS buffer plus 0.1% Tween 20) for 30 min at 60 rpm on rocker (LSE platform rocker, Corning) at room temperature. Next, blots were incubated with Monoclonal mouse α-human-CMAS 1° antibody (1:1000 in TBST with 10% BSA, cat. #: HPA039905, Atlas Antibodies). Note, the CMAS antibody was validated using SMART-pool siRNA (Horizon Discovery, **Figs. S5F & S8D**). After an overnight incubation at 4℃, blots were washed 2 × 30 seconds (sec.) with 0.1% TBST buffer. A secondary antibody was then added (α-rabbit IgG-HRP, 1: 6,000 in TBST with 10% BSA and incubated for 1 h at room temperature with shaking (60 rpm). Blots were then washed 2 × 30 sec. with 0.1% TBST buffer. Blots were developed using Clarity and Clarity Max Western ECL substrate according to the manufacturer’s instructions (Bio-Rad). Membranes were imaged in chemiluminescent mode (Azure 600, Azure Biosystems Inc.). All analysis was done in a minimum of biological triplicate.

Western blots were quantified using ImageJ software (ImageJ 1.54g, Java 1.8.0_345) (55). For each lane, signal was normalized to the Ponceau for that lane. For each blot the Ponceau normalized signal for miRNA mimics was divided by the NTC to give the normalized signal shown in all graphs. We tested for statistical significance using two different statistical tests: the one-sample *t*-test against NTC=1 and the paired *t*-test comparing Ponceau normalized NTC to miR for each blot. Both are shown in **Table S1**.

### Endogenous miRNA activity validation

miRIDIAN microRNA Hairpin Inhibitors (anti-up-miRs: -31-3p, -301b-3p, -941) and miRIDIAN microRNA Hairpin Inhibitor Negative Control (α-NTC, cat.#: IN-001005-01) were purchased from Dharmacon (Horizon Discovery, Cambridge, UK). The selected anti-miRs for CMAS protein were tested in MDA-MB-231 cell line. Cells were seeded and incubated as described for Western blot analysis. Cells were transfected with anti-miR (50 nM, using Lipofectamine™ 2000 transfection reagent in OptiMEM following the manufacturer’s instructions (Life Technologies)). After 12 h, media was changed to standard culture media. Cells were lysed 48 h post-transfection and analyzed for CMAS protein levels as previously described. All analysis was done in biological triplicate.

### Fluorescence microscopy

Cells were seeded onto sterile 22 × 22 glass coverslips placed into 35 mm dishes at a density of 2 × 10^5^ cells/ml in the appropriate media for the cell line as previously mentioned. After 24h, cells were transfected with miRNA mimics as previously described (see experimental procedure for “Western blot” section). At 48 h post-transfection, cells were washed with PBS (3 × 2 mL) and fixed with 4 % paraformaldehyde for 20 min. Cells were again washed with PBS (3 × 2 mL) and incubated with 1% BSA in PBS for 1 hour in the incubator (37℃, 5% CO_2_). The block buffer was removed and cells were incubated with Cy3-SNA (SKU #: CL-1303-1, Vector Laboratories; 1:300 (vol: vol), 2 mL total volume in PBS) for 1 hour in the incubator (37℃, 5% CO_2_). Coverslips were then washed (2 mL PBS, 3 x 4 min), and cells were counterstained with Hoechst 33342 (2 mL, 1 μg/mL in PBS, 15 min in incubator). The coverslips were mounted onto slides with 60 μl of mounting media (90% glycerol in PBS) and imaged with a Zeiss fluorescent microscope (Camera: Axiocam 305 mono, software: ZEN 3.2 pro). Specificity of SNA staining was confirmed using Neuraminidase A (expressed using NeuA construct, a gift from M.S. Macauley, University of Alberta) prior to Cy3-SNA staining. All analysis was done in biological triplicate.

### Flow cytometry analysis

miRNA transfection of samples for flow cytometry analysis was done as previously described. After 48 hours post-transfection, samples were trypsin digested (100 µl of 0.25 % trypsin per well in 6-well plate format). Up to 1 mL of 1X HBSS was added to remove all the cells from the flask, and cells were pelleted by centrifugation at 350 x g for 6 min. Cell pellet was resuspended in 1X TBS buffer containing 0.1% BSA and were counted using the cell counter. 100 µl of 5 x 10^5^ cells was counted per sample. As a negative control for lectin staining, untransfected cells were treated with neuraminidase (NEB, catalog #: P0720L) prior to staining. For staining, 15 µg/ml of FITC-SNA (Vector Labs, cat. #: FL-1302-2) in 1X TBS buffer was added onto each sample and incubated for 25 min at room temperature in the dark. Cells were pelleted by centrifugation at 350 x g for 5 min. Samples were washed with 1X TBS, 0.1% BSA 2x, and centrifuged at 350 x g for 5 min. Samples were resuspended in 400 µl FACS buffer (PBS, 0.1% BSA, 0.1% EDTA, 5 mM) and analyzed.

### Mutagenesis of miRNA sites on pFmiR-CMAS

The CMAS 3’UTR interactions with select miRNAs (down-miRs: -200c-5p, -550a-3p; up-miRs: -31-3p, -301b-3p) were analyzed using either Targetscan (20) or the RNAhybrid tool (40), which calculates a minimal free energy hybridization of target RNA sequence and miRNA. For down-miRs, the miRNA sites predicted by Targetscan were mutated. For up-miRs, the most stable predicted miRNA: mRNA interaction sites by RNAhybrid were selected. All sites were mutated to the corresponding miRNA sequence. Multiple mutation sites were designed, mutant primers were designed using NEB Base Changer version 1 (https://nebasechangerv1.neb.com) and ordered for synthesis by Integrate DNA Technologies (IDT). Primers are listed in **Table S2**. Multiple mutation sites were achieved using the Q5^®^ Site-Directed Mutagenesis kit (NEB, catalog #: E0554S) according to their protocols. Amplicons were cleaned up using Monarch PCR & DNA cleanup kit (catalog #: T1030S, NEB). Sequences for the mutant pFmiR-CMAS sensors were verified by sequencing and used in the miRFluR assay as described previously. A minimum of 5-wells were transfected per sensor and the analysis was done in three biological experiments. Data was normalized to the median control NTC (miR-625-5p) used with each sensor.

### Data availability

The authors declare that all data can be found in this document and its supporting files.

## Supporting information

This article contains supporting information:

Figs. S1-S8

Tables S1-S2

Datasets S1-S2 (separate excel files)

## Acknowledgements

We thank Dr. Matthew S. Macauley for generously providing the Neuraminidase (NeuA) plasmid. We thank Dr. Dawn Macdonald for providing comments on this manuscript.

## Author contributions

Conceptualization: F.J.C., L.K.M.; Methodology: F.J.C., L.K.M.; Investigation: F.J.C., J.N.R.; Validation: F.J.C., J.N.R., T.U.A.; Formal Analysis: F.J.C., J.N.R., L.K.M.; Visualization: F.J.C., J.N.R., L.K.M.; Writing-Original Draft: F.J.C., L.K.M; Supervision: L.K.M.

## Funding and additional information

Funding for L.K.M. comes from the Canada Excellence Research Chairs Program (CERC in Glycomics). Flow cytometry experiments were performed at the University of Alberta Faculty of Medicine & Dentistry Flow Cytometry Facility, RRID:SCR_019195, which receives financial support from the Faculty of Medicine & Dentistry and Canada Foundation for Innovation (CFI) awards to contributing investigators. Figs. 1, 2, 4, 6 were partially made using BioRender.

## Abbreviations

The abbreviation used are as follows: CMP: cytidine 5’-mono-phosphates; DNA: deoxyribonucleic acid; CMAS: cytidine monophosphate N-acetylneuraminic acid synthetase; ST6GAL1: α-2,6-sialyltransferase1; AGO2: Argonaute 2; PDAC, PAAD: pancreatic ductal adenocarcinoma; 3’UTR: 3’-untranslated regions; mRNA: messenger ribonucleic acid; microRNA, miRNA, miR: micro ribonucleic acid; siRNA: small interfering RNA; NTP: non-targeting control pool; NTC: non-targeting control, down-miR: downregulatory miRNA; up-miR: upregulatory miRNA; CD98hc, SLC3A2: neutral amino acid transporter; HRP: horseradish peroxidase; FITC: fluorescein isothiocyanate; Cy3: Cyanine3; SNA: *Sambucus nigra* agglutinin; kDa: kilodalton; WT: wildtype; MUT: mutant. IgG: immunoglobulin G; SDS-PAGE: sodium dodecyl sulfate polyacrylamide gel electrophoresis; PCR: polymerase chain reaction.

