## Supplemental Figs. S1-S8, Tables S1-S2 for "Profiling the Regulatory Landscape of Sialylation through miRNA Targeting of CMP- Sialic Acid Synthetase"

**This file contains:**

**Figs. S1-S8**

**Tables S1-S2**

**A**

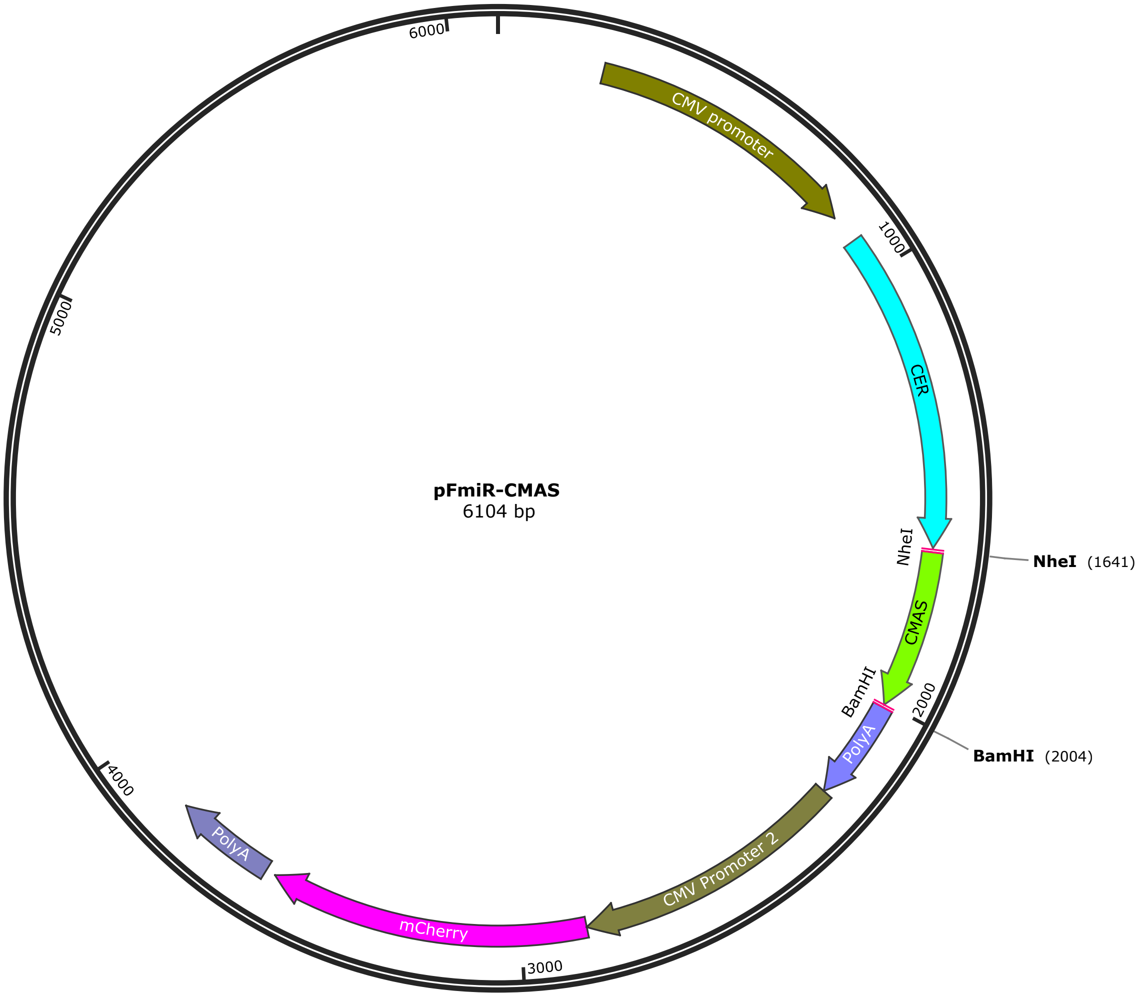


**B**

5’-AAATTAGCGTAATATTGAGAAAAAAATGATACAGCCTTCTTCAGCCAGTTTGCTTTTATTTTTGAT
TAAGTAAATTCCATGTTGTAATGTTACAGAGAGTGTGATTTGGTTTGTGATATATATATATTGTGCTCT
ACTTTTCTCTTTACGCAAGATAATTATTTAGAGACTGATTACAGTCTTTCTCAGATTTTTAGTAAATGC
AAGTAAGAACATCATCAAAGTTCACTTTGTATTGTACCCTGTAAAACTGTGTGTTTGTGTGCTTTCAAA
GATGTTGGGATTTTATTTATCTGGGGACAGTGTGTATGGTAAGACATGCCCTTCTATTAATAAAACT
ACATTTCTCAAACTTGA-3’

**Figure S1. pFmiR-CMAS map and sequence.** (A) pFmiR-CMAS plasmid map. (B) CMAS 3'UTR sequence. Sequence contains 357 base pairs.

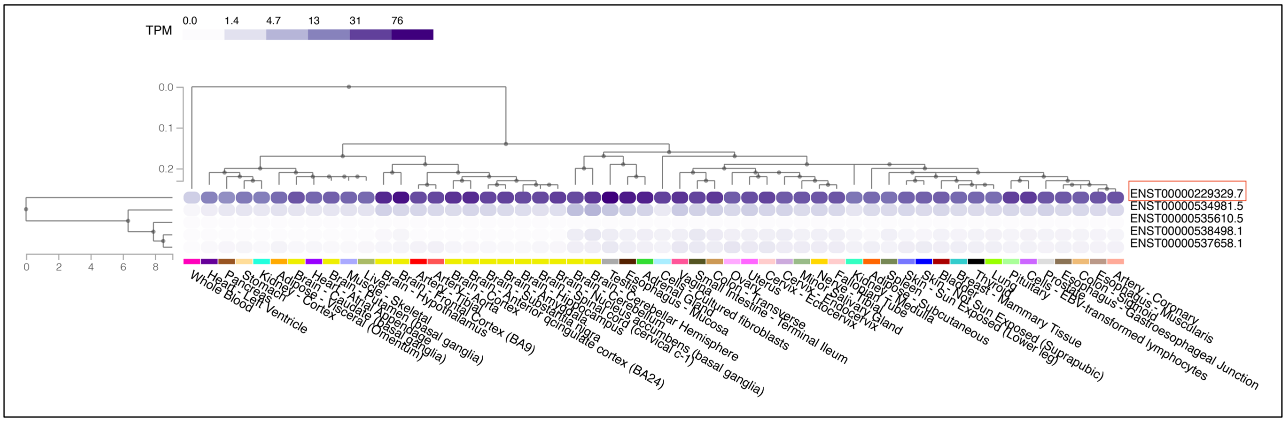


**Figure S2. Transcript expression level of CMAS in different tissues.** Transcript expression map is from GTex portal (https://www.gtexportal.org/home/transcriptPage for CMAS, downloaded 10/2024).

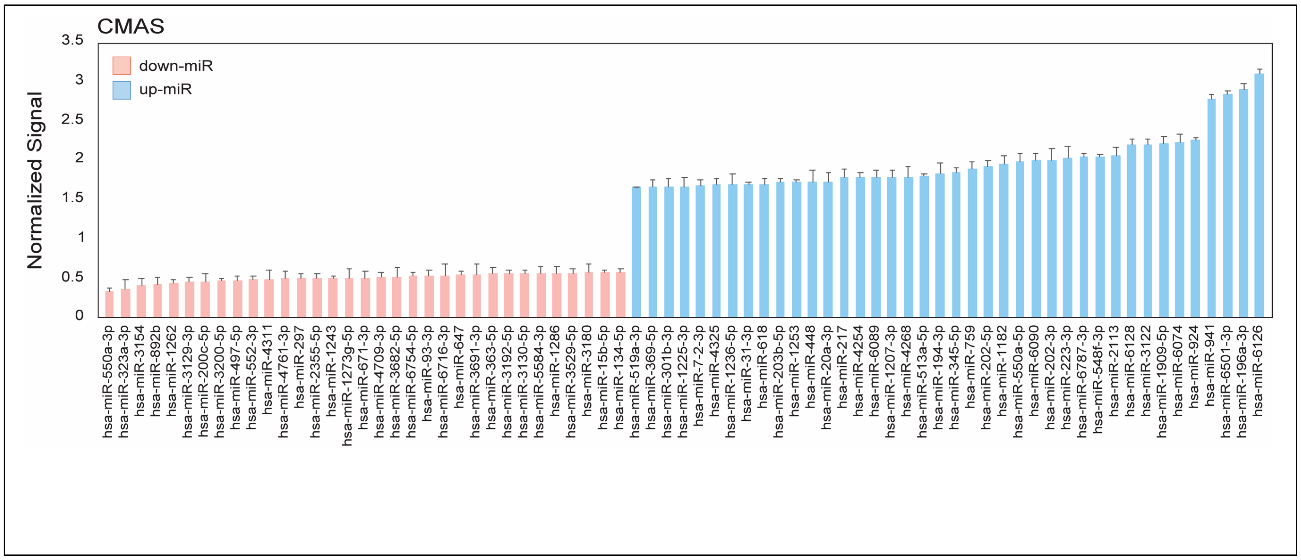


**Figure S3. miRFluR assay hits for CMAS.** (A) Bar graph of miRNA hits from 95% confidence interval for CMAS. Data are normalized over median. Error bars represent standard deviation of technical replicates (n=3).

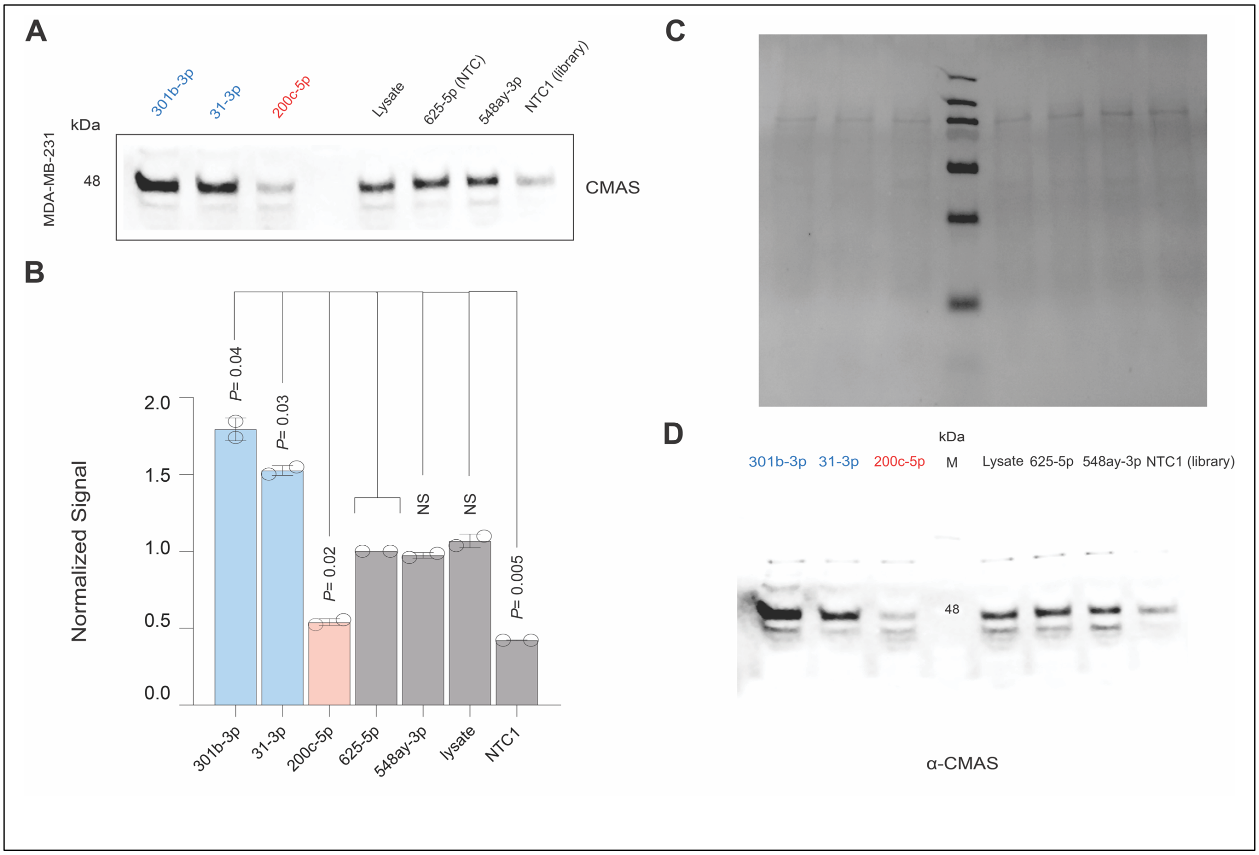


**Figure S4. Validation of miR-625-5p as new NTC.** (A) miRNA mimics of median controls (miRs: -625-5p (NTC), -548ay-3p -indicated on the blot) were transfected into MDA-MB-231 cells (50 nM miR, 48 h) prior to Western blot analysis. Representative data is shown. (B) Bar graph shows quantification of duplicate Western blot results as obtained in A. Signals were normalized to total protein on Ponceau. (C) Ponceau of Western blot shown in A. (D) Complete blot of Western shown in A, median control miRNAs are indicated.

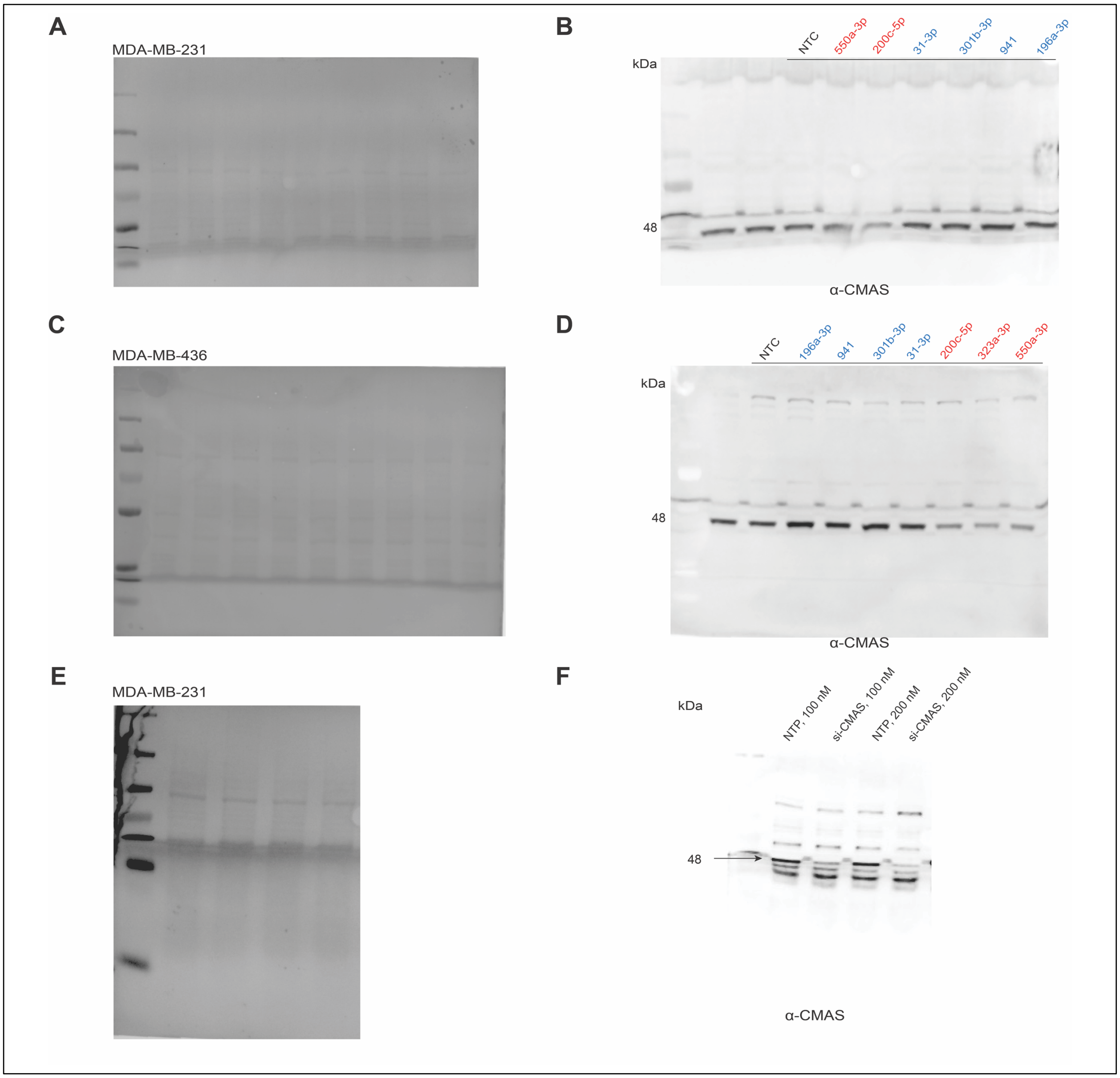


**Figure S5. Ponceau and whole Western blots for data shown in Fig. 3A and 3C.** (A, C) Ponceau staining of blots used in Fig. 3A, 3C (A: MDA-MB-231; C: MDA-MB-436). (B, D) Whole Western blot for data shown in Fig. 3A, 3C (B: MDA-MB-231; D: MDA-MB-436). (E, F) anti-CMAS antibody validation. (E) Ponceau staining of blot shown in F in MDA-MB-231. (F) Whole Western blot of siRNA validation of CMAS antibody.

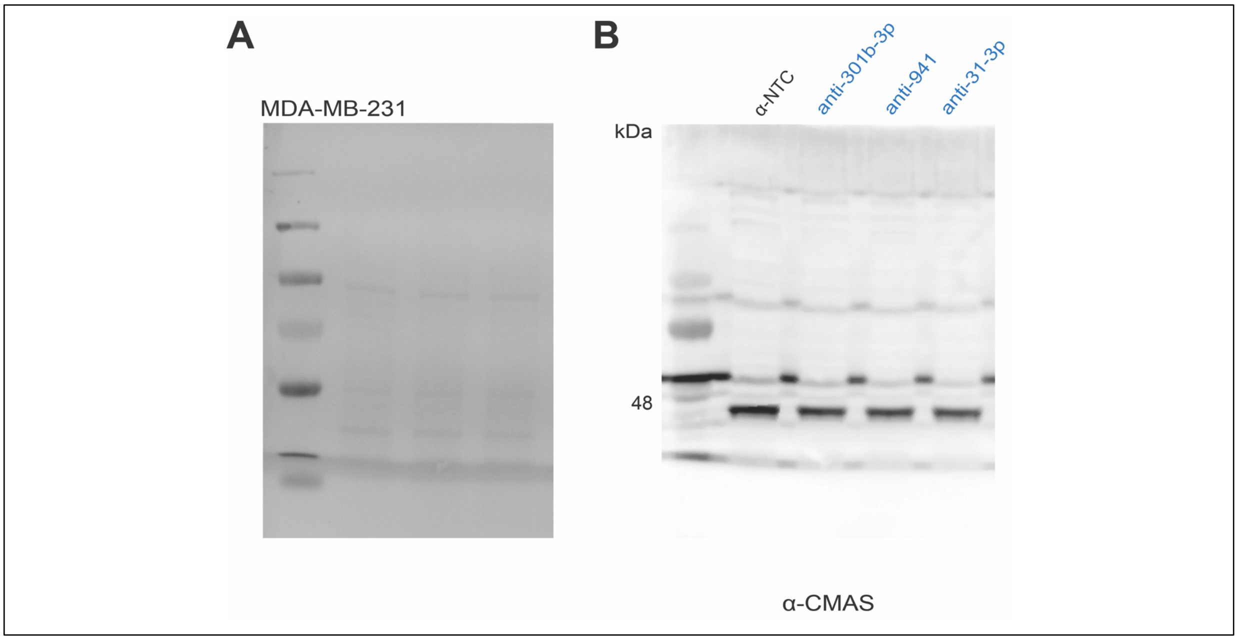


**Figure S6**. **Ponceau and Western corresponding to Fig. 4B in MDA-MB-231 cells**. (A) Ponceau staining of blot used in Fig. 4B. (B) Whole Western blot for data shown in Fig. 4B.

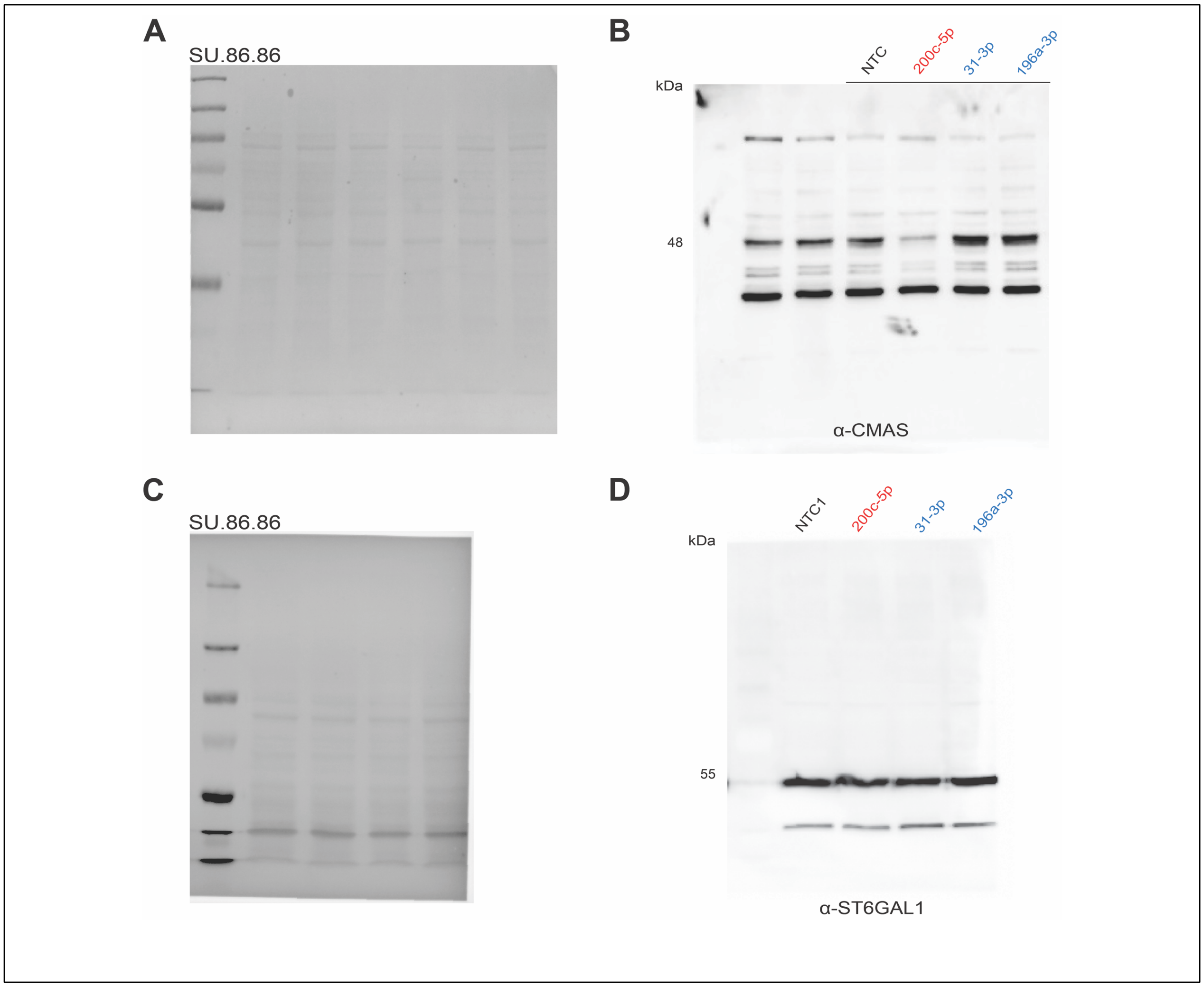


**Figure S7. Ponceau and Western corresponding to Fig. 6A and 6D in SU.86.86 cells**. (A, C) Ponceau staining of blots used in Fig. 6A, 6D (A: CMAS; C: ST6GAL1). (B, D) Whole Western blot for data shown in Fig. 6A, 6D (B: CMAS; D: ST6GAL1).

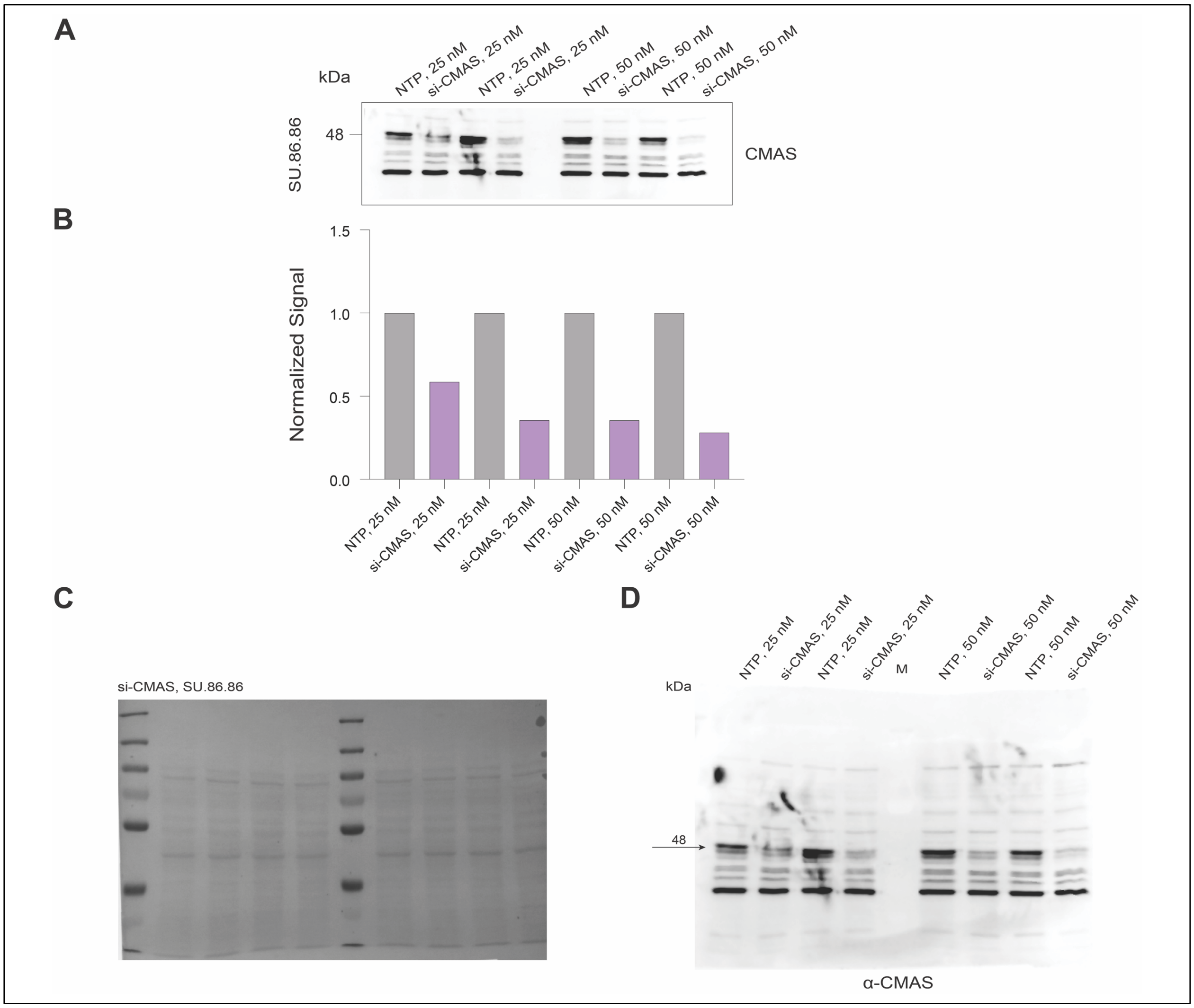


**Figure S8. Validation of antibody for CMAS in SU.86.86 cells.** (A) Western blot analysis of SU.86.86 cells treated with non-targeting pooled (NTP) or pooled siRNA targeting CMAS (si-CMAS). Cells were transfected into with varying amounts of siRNA (25, 50 nM) for 48 h prior to Western blot analysis. (B) Bar graph shows quantification of results shown in A. Signals were normalized to total protein on Ponceau. (C) Ponceau of Western blot shown in A. (D) Complete blot of Western shown in A.

**Table S1.** Statistical significance for Western blot experiments using both one-sample *t*-test and paired Student *t*-test.

| miRNA/ anti-miR | one-sample *t*-test | paired *t*-test | cell line |
|---|---|---|---|
| Fig. 3B | | | |
| up-miR | | | MDA-MB-231 |
| 196a-3p | 0.1 (NS) | 0.2 (NS) | |
| 941 | 0.02 | 0.1 (NS) | |
| 301b-3p | 0.0004 | 0.03 | |
| 31-3p | 0.01 | 0.0003 | |
| down-miR | | | |
| 200c-5p | 0.003 | 0.007 | |
| 323a-3p | 0.03 | 0.04 | |
| 550a-3p | 0.002 | 0.005 | |
| Fig. 3D | | | |
| up-miR | | | MDA-MB-436 |
| 196a-3p | 0.01 | 0.08 (NS) | |
| 941 | 0.01 | 0.03 | |
| 301b-3p | 0.0005 | 0.04 | |
| 31-3p | 0.01 | 0.03 | |
| down-miR | | | |
| 200c-5p | 0.004 | 0.009 | |
| 323a-3p | 0.01 | 0.04 | |
| 550a-3p | 0.02 | 0.05 | |
| Fig. 4C | | | |
| anti-up-miR | | | MDA-MB-231 |
| anti-301b-3p | 0.009 | 0.04 | |
| anti-31-3p | 0.02 | 0.05 | |
| anti-941 | 0.003 | 0.03 | |
| Fig. 6B | | | |
| up-miR | | | SU.86.86 |
| 196a-3p | 0.004 | 0.02 | |
| 31-3p | 0.0007 | 0.02 | |
| down-miR | | | |
| 200c-5p | 0.02 | 0.003 | |

**Table S2.** Primer sequences for PCR amplification of wild-type (WT) 3'UTR (A) and for site directed mutagenesis of CMAS (B).

| Primer Name | Sequence (5' → 3') | Sample |
|---|---|---|
| (A). PCR amplification of CMAS 3'UTR | | |
| CMAS-FWD[a] | AGTAAATGCAAGTAAGAACATCATCAAAG | gDNA, HEK-293T |
| CMAS-REV[a] | CCCCAGATAAATAAAATCCCAACAT | gDNA, HEK-293T |
| (B). PCR amplification of CMAS mutant 3'UTRs | | |
| 200c-5p-CMAS-MUT-FWD | atgcattctgtTGCCCTTCTATTAATAAAAC | pFmiR-CMAS |
| 200c-5p-CMAS-MUT-REV | acacactgtcccCAGATAAATAAAATCCCAACATC | pFmiR-CMAS |
| 550a-3p-CMAS-MUT-FWD | atgcattctgtTGCCCTTCTATTAATAAAAC | pFmiR-CMAS |
| 550a-3p-CMAS-MUT-REV | acacactgtcccCAGATAAATAAAATCCCAACATC | pFmiR-CMAS |
| 31-3p-CMAS-MUT-FWD | aaccattcagttGCCCTTCTATTAATAAAACTAC | pFmiR-CMAS |
| 31-3p-CMAS-MUT-REV | tgtgtgactggcCAGATAAATAAAATCCCAACATC | pFmiR-CMAS |
| 301b-3p-CMAS-MUT-FWD | ataaataacacgacTACTTTTCTCTTTACGCAAG | pFmiR-CMAS |
| 301b-3p-CMAS-MUT-REV | aataacagtttcgaAATCACACTCTCTGTAAC | pFmiR-CMAS |

[a] FWD, forward; REV, reverse.
